# Development of a displacement-controlled uniaxial-strain bioreactor for high-throughput, dynamic *in vitro* cell culture

**DOI:** 10.64898/2025.12.18.695040

**Authors:** ML Eames, E Pickering, P Subedi, MA Woodruff, TJ Klein

## Abstract

Bone is a dynamic tissue that experiences a wide range of forces during regular daily locomotion. This environment of dynamic strain strongly influences the architecture of the extracellular matrix, and it can impact the rate that bone adapts or recovers after an injury [1], [2]. Cell research is commonly performed in mechanically static conditions in the base of well plates, yet this is a far cry from the conditions natural to osteoblasts. To better understand the behaviours of osteoblasts, it is important to ensure that platforms are available to perform cell culture experiments in mechanically dynamic environments [3]–[5]. For this reason, we have designed a bioreactor system that imparts tensile strain onto flat, cell-seeded scaffold constructs *in vivo* within custom well plates. The bioreactor system has 36 separated wells spread across four mechanical actuation units and can apply up to 30% tensile strain to a 15 by 9 mm area of the constructs, operating within an incubator. The wider body of mechanostimulation research also shows that different many cell types, from neurons [6] to cardiac tissue [7], [8] and more [2], [6], [9]–[11], can be stimulated with a range of stimulus and have a wide array of responses, and it is expected that the bioreactor will also be useful for this research. To be accessible to research groups, the bioreactor has predominantly been constructed from widely available components and materials. Most parts fabricated using 3D printing, and all of the electronics can be found within a typical 3D printer DIY assembly kit. The device was validated for use in a 28 day in-vitro dynamic culture of osteoblasts on melt-electrowritten polycaprolactone scaffolds. The cell constructs were strained to 4% at 0.5 Hz on days 25-27 and removed on the 28^th^ day. Cells were observed to elongate and align within some regions of high local strain following the 3 days of stimulation.

## 2 Introduction

Throughout regular, daily locomotion and activities the human skeleton experiences a range of mechanical forces [12], [13]. These forces play an integral role in the development of the skeletal system by guiding the formation and remodelling of bone tissue at a cellular level. Through mechanotranduction, osteocytes sense the types, magnitudes and directions of strains and influence the balance between osteoblast-driven matrix mineralisation and osteoclast-driven resorption [14], [15]. An accurate understanding of how bone tissue develops is vitally important within the context of wound healing, as it can lead to more effective bone graft treatments in particular for critical-sized defects [16], [17].

Mechanoculture experiments culture cells under mechanically dynamic environments to study their behaviour and responses directly to and throughout the stimulus [18]–[22]. This research requires the use of bioreactors to impart mechanical loads onto cell populations. Some of the earliest and simplest forms of dynamic bioreactors for cell culture research were the rotating wall vessels and spinner flasks, tissue culture flasks that would cause medium to flow over cells by their continual rolling or through the use of a stirrer bar at the base [4], [23], [24]. More recently, tension, compression, and shear have been applied independently or in combination and found to promote cell proliferation and matrix deposition, and have also been observed to have differential effects on cell cultures [25], [26]. Cyclic compression and hydrostatic pressure have been found to upregulate chondrogenesis of human mesenchymal stem cells (MSCs) in scaffolds, particularly boosting the production of the collagen II protein associated with articular cartilage alongside increasing MSC proliferation [27]–[30]. Cyclic tensile strain (CTS) however has been found to preferentially upregulate osteoblast proliferation, mineralisation, and ALP activity while suppressing chondrogenesis [1], [31]–[36]. CTS can specifically promote the production of the collagen-I and collagen-III proteins associated with native bone tissue without also increasing the production of collagen-II in opposition to the effects of cyclic compressive strain (CCS) [37]–[39]. This suggests CTS is more suitable for promoting osteogenesis within bone cell constructs, though compared to research into the effects of compression or perfusion on 3D constructs this is scarcely investigated.

A range of bioreactors for applying fluid or compressive shear to many samples simultaneously have been proposed within the literature, and several have been developed into commercial products [2], [4], [18]–[20], [27]–[30], [40]–[50]. As a result, research into the effects of these strain types is well established. Research into the effects of tensile strain on 3D cell constructs is however addled by a lack of high throughput devices appropriate for rigorous mechanoculture experiments. Commercial platforms such as by CellScale, Ebers and Flexcell do exist and have been used in various studies [31], [39], [51]–[54], however their usage is quite limited, especially for stimulating 3D cell constructs. Many research groups to develop their own bioreactors internally, dedicated to their own research [1], [5], [55]–[68]. These bespoke devices have enabled individual groups to uncover new understanding on how mechanical conditions influence cell development and behaviour, but these results are poorly replicable as they are strongly tied to the custom devices, which are rarely described. This frustrates the development of clear understandings of how mechanical conditions affect cell behaviour.

To support more robust progression of mechanostimulation research for osteogenic CTS, we have developed a low-cost, open-source bioreactor, the ‘OpenStrain Bioreactor,’ to stimulate cell constructs with mechanical strain through controlled displacement. The device requires little mechanical or programming expertise to assemble and use, and allows for higher-throughput experiments than similar available platforms.

## 3 Materials and methods

### 3.1 Bioreactor design and fabrication

#### 3.1.1 Design methodology and criteria

The OpenStrain Bioreactor system was developed over ∼24 months using an iterative design process, in which design generations were produced and evaluated with respect to design criteria seen in **Table 1**, then adapted to more closely align with the criteria and user needs. The design criteria were developed based on the analysis of existing mechanical strain bioreactors and ensure the device would be an effective tool for efficient mechanostimulation research. Solidworks 2022 (Dassault Systems, SA, France) was used to develop computer-aided design (CAD) models. Many versions of the bioreactor were produced as the design was refined up until the last, seen on the far right of **1Figure 1**

**Figure 1.**
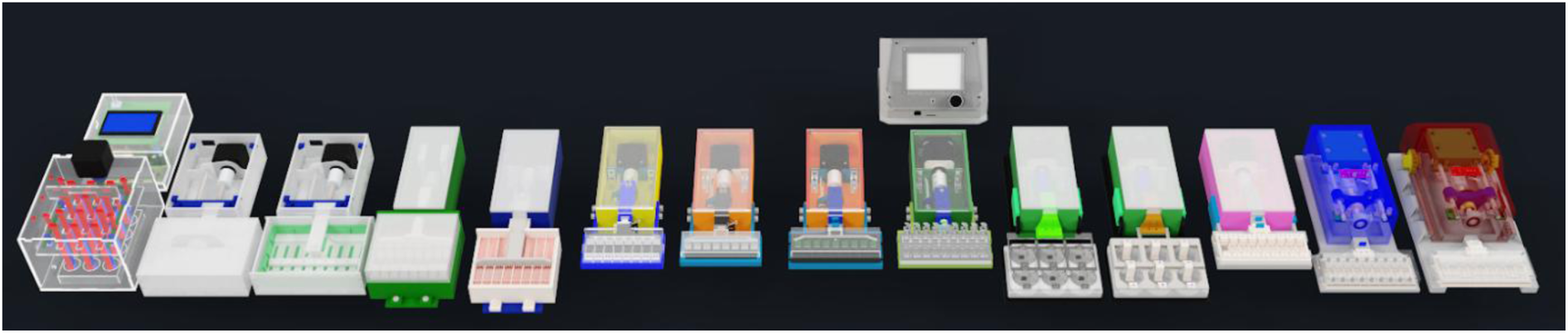
Prototype bioreactor designs produced through the iterative design process, progressing from earliest on the far left to project-final (the OpenStrain Bioreactor) on the far right.

**Table 1.** Design criteria for the development of the bioreactor system.

| Critical design criteria |
| --- |
| The device will reversibly hold cell constructs within a water-tight volume that can be sealed as securely as a standard polystyrene cell culture plate. |
| The device will apply cyclic tensile strain of up to at least 30%. |
| The device will operate within an incubator for at least three days following appropriate sterilisation by steam autoclave or ethanol evaporation. |
| The device will be reusable, using materials and parts that can be reliably and efficiently sterilised or else inexpensively refabricated/purchased. |
| Preferable design criteria |
| It will be possible to purchase or fabricate all components of the device for under AUD \$2000. |
| It will be possible to assemble the device without advanced technical skill beyond what would be required to assemble a commercial 3D printer kit. |
| The device will maximise experiment throughput by stimulating many samples simultaneously and minimising required media volumes. |
| It will be possible to extract samples for analyses without releasing all samples simultaneously. |
| The device will have small footprint to allow multiple to operate within a single incubator. |

#### 3.1.2 Bioreactor system overview

The ‘OpenStrain Bioreactor’ is a device for applying cyclic strain to flat, rectangular tissue-engineered constructs within an incubator for mechanically dynamic cell culture (**Figure 2**). The system consists of a ‘Control Unit’ and individual ‘Bioreactors’. The control unit can run up to three bioreactors simultaneously with a dedicated limit switch internal to each bioreactor, with the capacity for expansion to up to five bioreactors pending advanced firmware updates. The control unit is connected to each bioreactor via a flat ∼1.5 m umbilical cable that can pass through the door of an incubator without interfering with the seal, so that the user can operate the control unit from outside the incubator while it controls bioreactors within.

**Figure 2.**
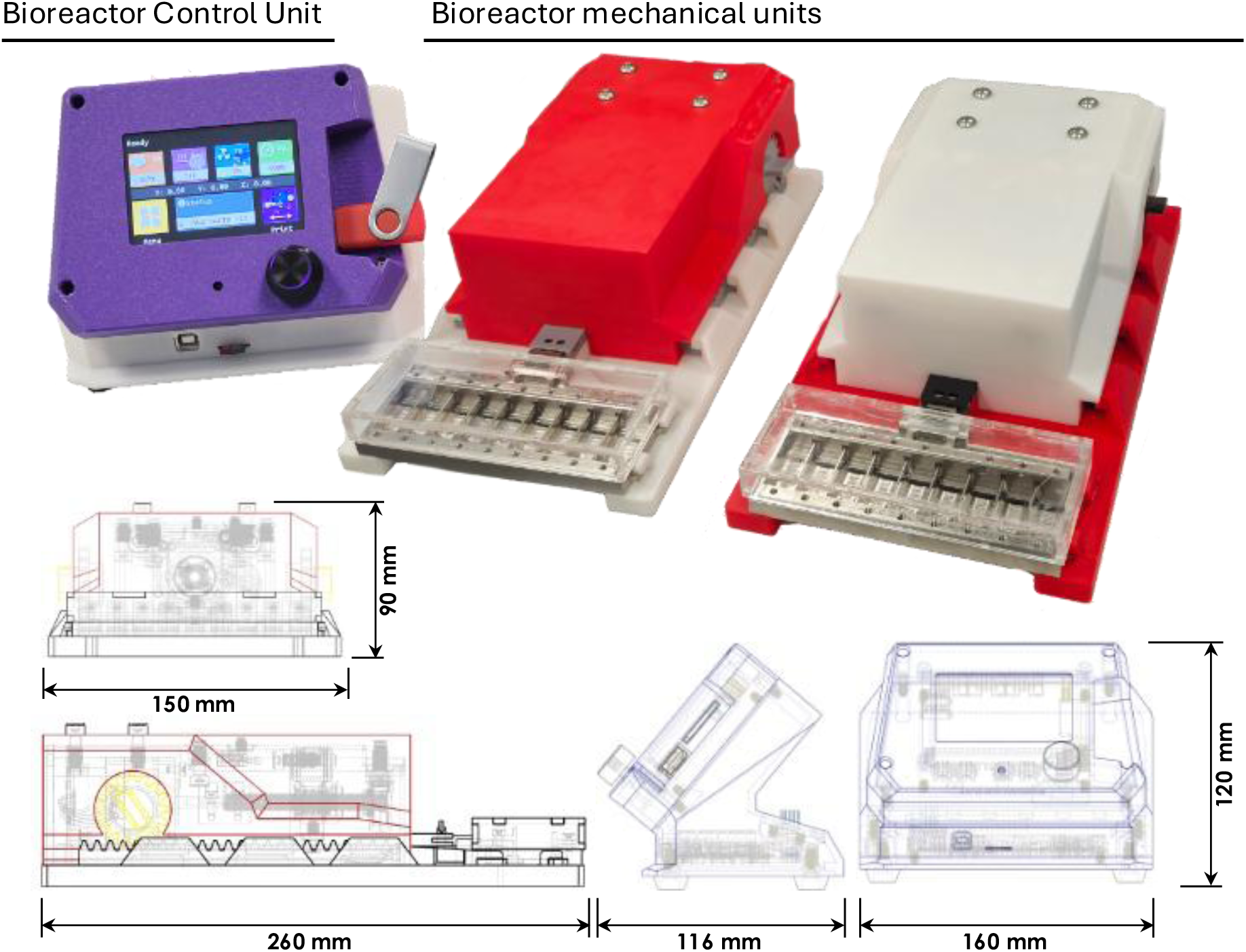
Summary of OpenStrain Bioreactor system and external dimensions. Bioreactor-to-control unit communication and power cables and the control unit power cable omitted for brevity.

#### 3.1.3 Electro-mechanical system

The control unit uses a BigTreeTech SKR 3 3D printer control circuit board installed with a customised version of Marlin 2.1.2.1 3D printer firmware (Available on github: https://github.com/teameames/OpenStrain-Bioreactor) to drive closed-loop stepper motors (BigTreeTech S24C v1.1) within each bioreactor and monitor signals from limit-switches within each bioreactor. Loading regimes are written as Gcode scripts using a custom MATLAB script based on inputs such as the required strain magnitude, rate, and duration. These can then be loaded onto the internal storage of the control unit using a MicroSD card or a USB and run independently of a computer. The progress of the loading regime is displayed on the LCD touch screen (BigTreeTech TFT 35) of the control unit and can be adjusted, paused or terminated from the display. The closed-loop motors self-correct errors due to half-steps and continually display the error of their position on individual displays visible through a clear acrylic panel on the back of each bioreactor.

The OpenStrain Bioreactor applies uniaxial strain using an ACME leadscrew mechanism, with a four-start 8 mm metric leadscrew interfacing with a Delrin anti-backlash nut fixed to a ‘lineariser’ moving carriage, for precise movement (**Figure 3**). The lineariser is attached to the pull arm which connects to the well plate assembly to transmit displacement from the actuator to samples within the well plate. The lineariser has an adjustable spring-loaded screw that activates a limit switch fixed within the housing of bioreactor. When the homing option is selected on the control unit display, the lineariser drives towards the limit switch to calibrate the mechanical assembly and define the zero point of the axis. The maximum displacement from this point is set within the firmware to prevent the lineariser from travelling beyond the range of the attached well plate.

**Figure 3.**
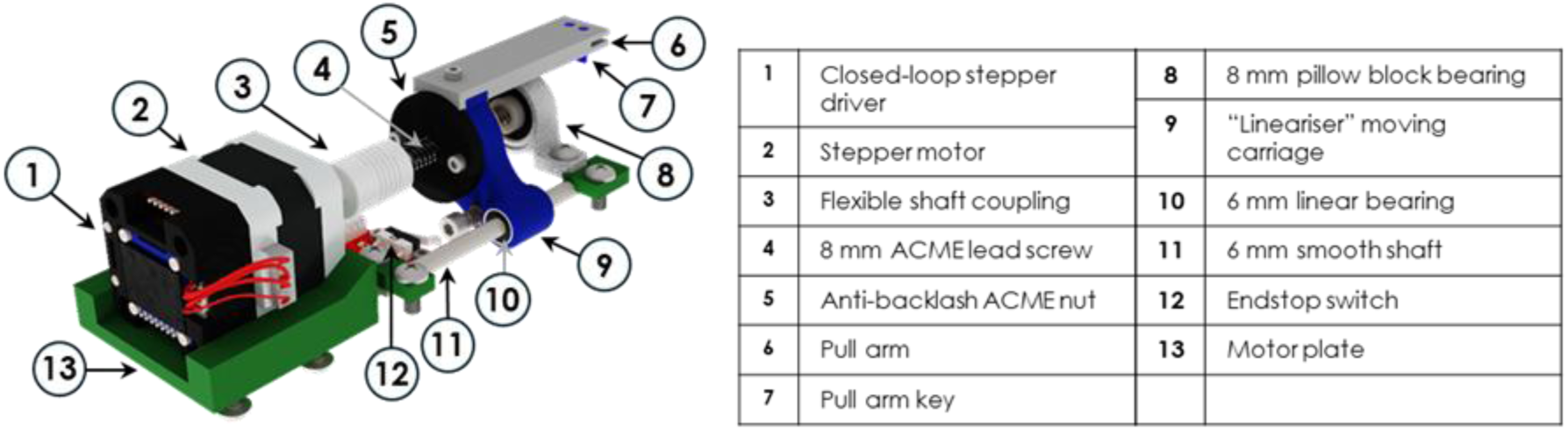
Detailed view of OpenStrain Bioreactor mechanical assembly with part identification.

Components of the mechanical assembly were purchased from Makerstore (Carrum Downs, Victoria), including the lead screws, smooth shafts, shaft couplings, bearings, and anti-backlash lead nuts. Electronic components were purchased from Biqu Equipment (Shenzhen, Guangdong, China), including the main control board, display, and closed loop motors.

#### 3.1.4 Well plate assembly

Each bioreactor unit stimulates one custom well plate of constructs (**Figure 4**). Each well plate has 9 separated wells that each receive the same applied displacement and sample strain, with the constructs held by screw-fastened clamps attached to the pull arm and to the opposite side of the well plate. The well plates support constructs between 22 and 24 mm long, and up to 10 mm wide, with a minimum separation of 15 mm between the clamps. The maximum stroke of the pull rake is 4.5 mm, equating to a maximum sample strain of 30% extending the free area of constructs from 15 mm to 19.5 mm.

**Figure 4.**
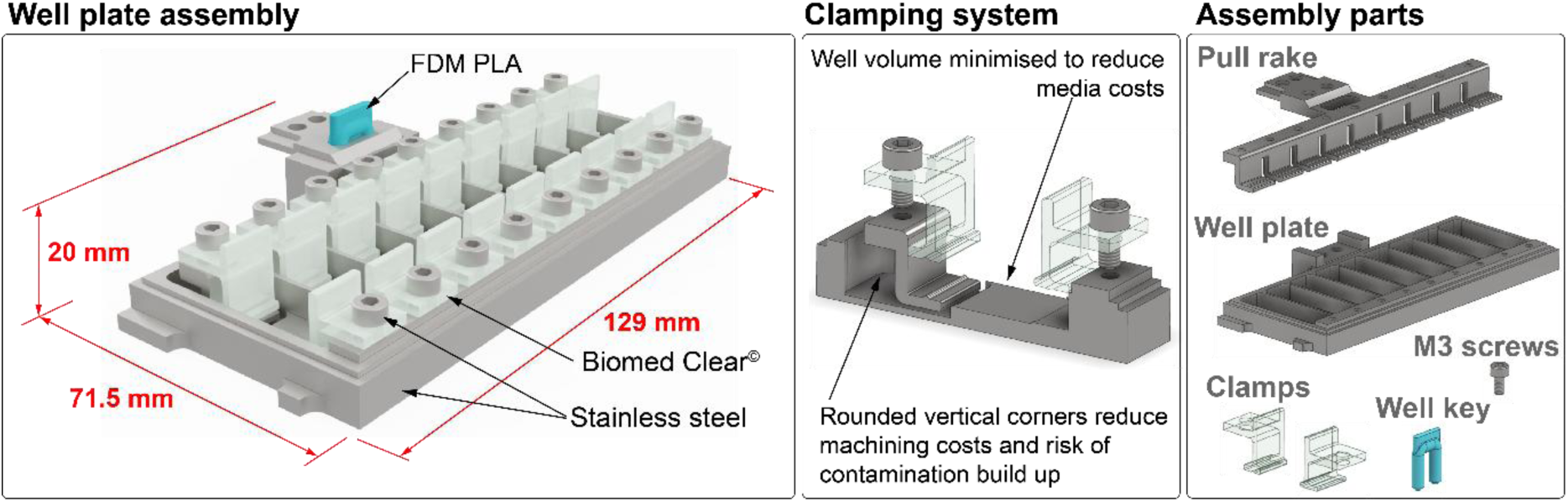
Sectioned view of stainless-steel well plate with screw-fastened Biomed Clear clamps. Clamps are 10 mm wide and approximately 13 to 15 mm high.

The clamps, screws, well plate and pull arm are all made from biocompatible and autoclavable materials so they can be sterilised and reused. The well plate and pull arm were produced by the QUT Design and Fabrication Research Facility (DeFab) by CNC machining from 316 stainless-steel. This was completed in-house at the university and through 3^rd^ party manufacturing houses such as SunPE, producing equivalent parts. The following design constraints were applied to reduce machining costs and complexity:

§ Minimum inner radius of 2 mm for inner corners axial to the milling bits.

§ Minimum inner radius of 0.1 mm for inner corners perpendicular to the milling bits.

§ Maximum depth of 15 mm for details requiring a 2mm radius bit.

The well plate was intentionally designed to be compatible with three-axis CNC mills, to prevent the need for repositioning the part or using a more expensive five-axis CNC mill. This could not be achieved for the design of the pull rake due to the necessity of overhangs in its design, so it would still require a five-axis CNC mill for fabrication or repositioning within a three-axis mill.

The clamps wer made from Formlabs BioMed Clear resin, printed on a Formlabs 3B+ resin printer (Formlabs, Somerville, Massachusetts). The screws were ‘off-the-shelf’ fasteners, made from 304 stainless-steel. A clear acrylic lid is used to protect the samples during experiments, cut from 3 mm acrylic sheet using a Trotec Speedy 300 precision laser cutter (Trotec Laser, Marchtrenk, Austria). Acrylic parts were assembled using cyanoacrylate superglue. 24 hours after gluing, the acrylic parts were soaked in 2 L of deionised water for 2 weeks, with water changes every 2 - 3 days, to prevent the glue from leeching and interfering with cell constructs.

The screws and clamps can each be removed individually, allowing for individual samples to be removed from the device at different timepoints during mechanically dynamic experiments using the device. Each well of the well plate requires 700 - 800 µm of media solution to ensure constructs are covered. Up to this maximum, cell media will not spread into adjacent wells or come into contract with screws or screw threads, which can be more difficult to thoroughly clean. The inner corners of the wells have fillet radii of at least 0.1 mm to reduce the likelihood of cells attaching.

#### 3.1.5 3D-printed control unit and bioreactor unit housings

The control unit and bioreactor units use FDM 3D printable housings that can be printed from PLA or other common FDM filaments (e.g. ABS, PETG) (**1Figure 5**). FDM parts were printed using generic PLA filament on a Prusa MK3S and a Prusa MK4 (Prusa Research, Prague, Czech Republic) and on a Creality K1 Max (Creality, China). Brass heat inserts installed into each part allow the housing to be assembled quickly with screws an enable hot swapping of parts and easy maintenance, if needed. A panel of clear acrylic is used for the rear panel of the bioreactor unit housing so the display on the stepper motor encoder can be viewed. Stainless steel screws are used to secure the mechanical assembly to the bioreactor unit housing to prevent corrosion from the high humidity within an incubator. The main housing of the bioreactor unit is secured to the base plate using a quickly fastened rack and pinion mechanism.

**Figure 5.**
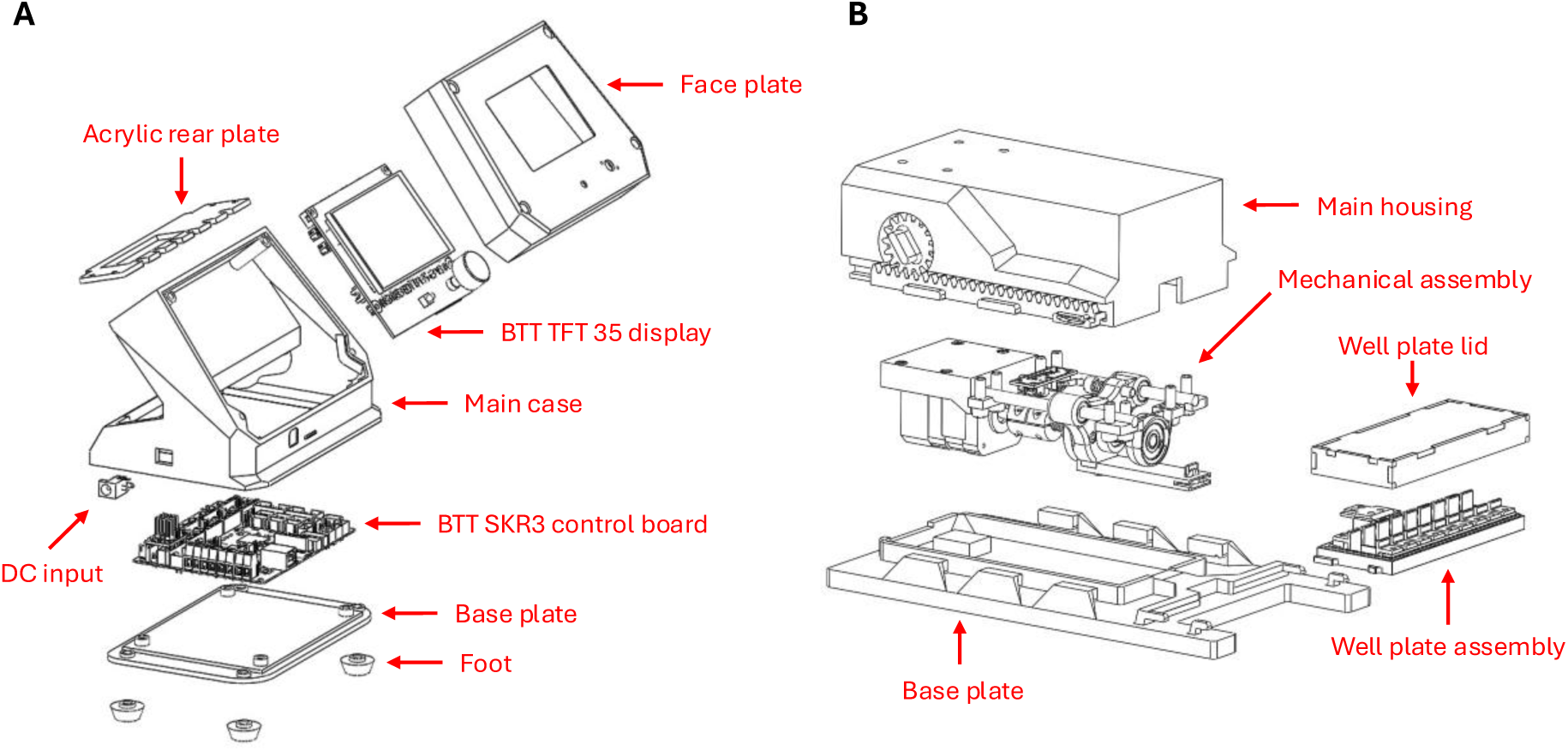
Exploded view of the **A:** control unit assembly and **B:** drive box assembly for the OpenStrain Bioreactor.

#### 3.1.6 Fixed microscopy under strain imaging tool

An imaging tool was designed within the OpenStrain bioreactor ecosystem to enable fixed samples to be imaged with microscopy under static strain (**Figure 6**). The device uses a crank-lever arm to slide a movable clamp, with sockets in the crank arm that produce sample strains of 0%, 4%, 5%, and 10% for 15 mm scaffolds equivalent to those used in the OpenStrain well plate.

**Figure 6.**
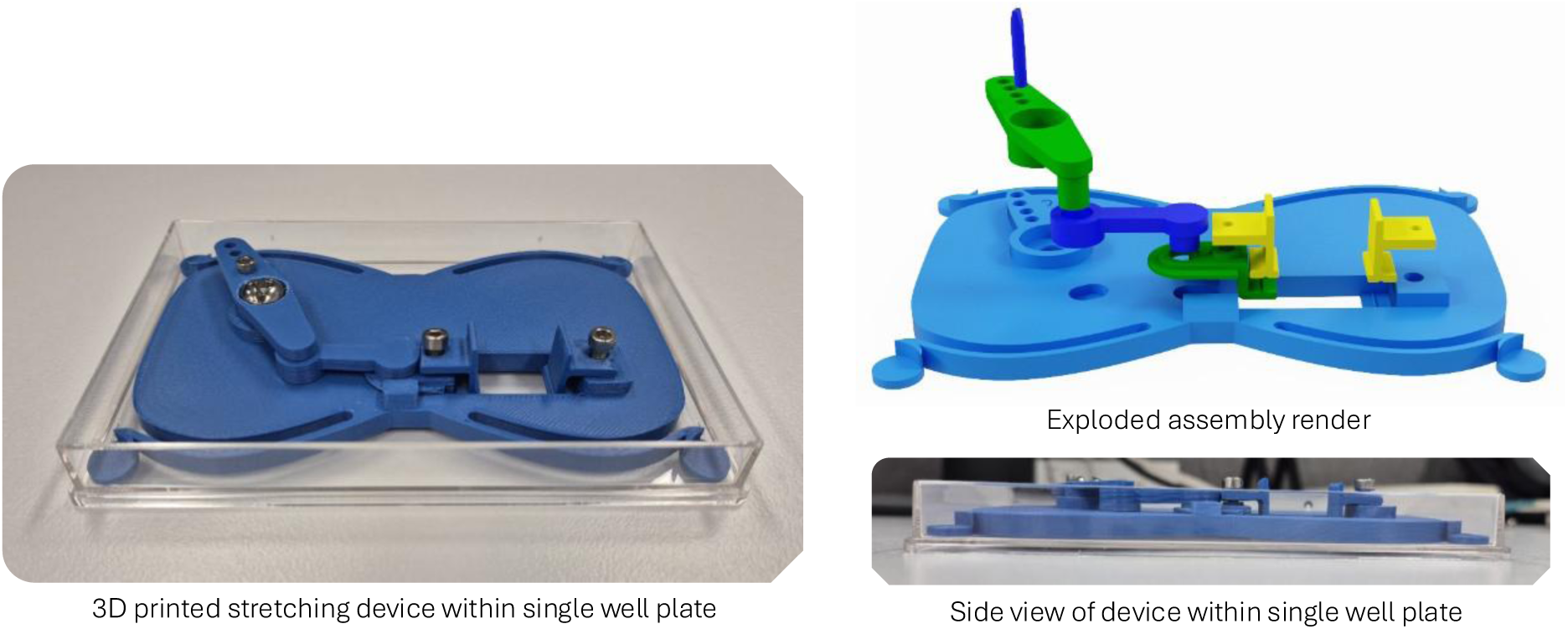
Device for applying static strain to fixed samples from the OpenStrain Bioreactor during microscopy. The minimum separation between the clamps is 15 mm. The maximum displacement of the moving clamp is 1.5 mm, equating to a maximum sample strain of 10%. The sockets on the crank arm (green in the assembly render) correspond to 0%, 4%, 5%, and 10% sample strain for a 15 mm scaffold.

**Figure 7.**
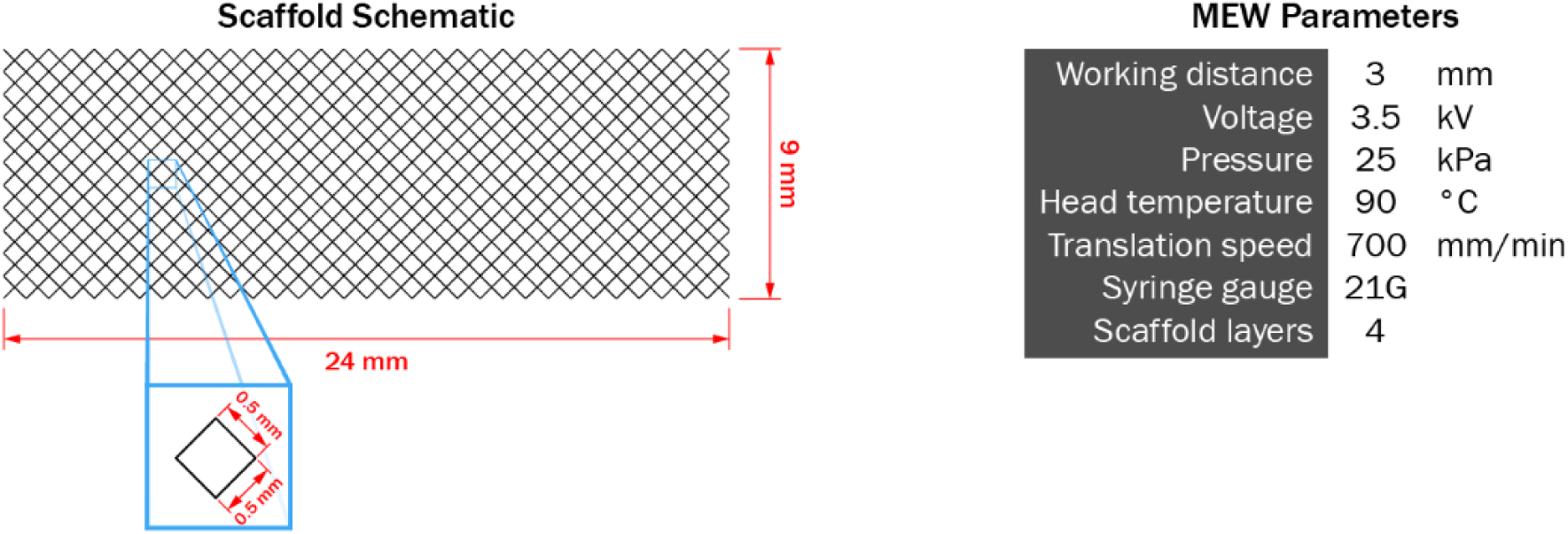
Scaffold schematic and MEW print parameters for dynamic study tissue constructs.

The clamping system used for the device is the same as OpenStrain well plate. The device is fabricated FDM printed PLA with stainless steel screws and nuts, brass inserts, and a polystyrene single well plate. The well plate enables images to be taken with sample within PBS and fits the plate holders of most fluorescence microscope stages. The assembly is 16.75 mm high including the blue socket peg, allowing it to fit within upright microscopes. Flexible feet at each corner of the device push outward on the inner walls of the well plate to maintain the position of the imaging window and fit within single well plates with varied internal dimensions.

### 3.2 Functional validation of the bioreactor system

#### 3.2.1 Validation of movement precision

The precision of the bioreactor systems was measured by running a cyclic program for up to 4 hours and videoing the movement of the tip of the pull arm. The videos were then processed in a custom MATLAB script that tracked the movement and reported the variance in the location of the pull arm at the 0 % strain point of the loading cycle. The script tracked movement using the MATLAB computer-vision toolbox to track dark dots drawn on the pull arm as visual landmarks, and by measuring the movement of regions of stark changes in contrast along a slice of pixels in the video in line with the direction of movement.

The accuracy and precision of the bioreactor pull arm moving to specified locations within its maximum movement were tested in accordance with the methods described by Janvier and Banik for similar custom tensile strain devices [5], [69]. Briefly, the pull arm was moved in 100 µm increments over 3000 µm using the on-board movement commands on the control unit. The position of the pull arm following each movement was measured using precision callipers. This experiment was performed in triplicate.

#### 3.2.2 Validation of thermal stability from stepper motors

The heat output of the motor during normal actuation was investigated using a FLIR T660 infrared camera (Teledyne FLIR, USA) to indicate if it would be likely to interfere with the automatic temperature regulation of a typical CO_2_ incubator. A four hour cyclic loading program typical of what would be used in dynamic cell culture was run on the bioreactor in ambient temperature, and the relative heat of the motor and drive box taken from a sampling point within the camera view finder.

#### 3.2.3 Static cytotoxicity study

A brief pilot study was conducted to confirm whether the custom-machined stainless-steel well plates were non-cytotoxic and would support *in vitro* cell culture of cell-seeded constructs. The study also briefly investigated the effects of different seeding methods on metabolic activity and cell attachment. The pilot study investigated four comparative groups displayed in **Table 2**.

**Table 2.** Comparative groups within pilot static cytotoxicity study.

| Sample Group ID: | SSIN | SSOUT | SBS200 | SBS100 |
| --- | --- | --- | --- | --- |
| Main culture well | Stainless-steel | Stainless-steel | SBS well | SBS well |
| Scaffold dimensions (mm) | 24 x 9 | 24 x 9 | 15 x 9 | 15 x 9 |
| Seeding volume ($\mu\text{L}$) | 200 | 200 | 200 | 100 |
| Seeding location | In culture well | In SBS well, moved to culture well after 2 hours | In culture well | In culture well |

The cytotoxicity of the stainless-steel well plate and pull arm were assessed in a 7-day cell culture by culturing PCL MEW scaffolds seeded with MC3T3-E1 murine pre-osteoblast cells within each of the wells. Scaffolds were printed using custom MEW printers [70]. Scaffolds within groups SSIN and SSOUT were printed with 24 x 9 mm outer dimensions to match the specifications of the stainless-steel well plate. Scaffolds within groups SBS200 and SBS100 were printed with 15 x 9 mm outer dimensions to simulate the free area of scaffolds within clamps when clamped in the stainless-steel plate. All scaffolds used 500 µm rectangular unit cells.

The hydrophilicity of the scaffolds was increased by etching through immersion in NaOH for 15 minutes, followed by 5 washes in excess deionised water. The scaffolds were sterilised by exposure to UV overnight in a biosafety cabinet prior to the experiment. Immediately prior to the experiment, the stainless-steel parts were cleaned with 70% ethanol and then sterilised by autoclave at 121 °C for 20 minutes and allowed to cool in sterile conditions. Metabolic activity of cells attached to scaffolds was assessed using AlamarBlue® assay reagent, in fresh wells to prevent contribution from cells attached to the wells. An ethidium homodimer-1 (Invitrogen) and Calcein-AM (Invitrogen) Live/Dead fluorescence assay was used to assess cytotoxicity at Day 3 and Day 7 using an AxioObserver Z1fluorescence microscope (Carl Zeiss, Germany).

#### 3.2.4 Dynamic *in vitro* tissue construct model

The effect of cyclic loading on MC3T3-E1 cell/PCL MEW scaffolds constructs was investigated using the device. PCL scaffolds were printed from 45K MW PCL (Sigma Aldrich) with a simple repeating diamond unit cell (Figure X). Fibres had an average diameter of 48 µm and 500 µm pore size. To increase hydrophilicity and promote even cell seeding, scaffolds were plasma treated with O_2_/Ar (15:5) for 2 minutes at 38 W in a vacuum plasma cleaner (PDC-002-HP, Harrick Plasma, USA). They were then sterilised by soaking in 85% ethanol for 30 minutes, dried by evaporation within a biosafety cabinet, and seeded with cell within 24 hours of plasma treatment.

Scaffolds were seeded with murine calvarial osteoblastic cells (MC3T3-E1) in 6-well plates at a concentration of 750,000 cells per mL, using two 100 µL aliquots per sample. The seeding solution was made up on α-MEM, 10% fetal bovine serum, and 1% penicillin-streptomycin (Thermo Fisher) media solution. The aliquots were deposited approximately in the centre of the left and right halves of the region of the scaffold that would be between the bioreactor clamps, as viewed in landscape. The seeding volume was deposited in two aliquots to encourage even cell dispersion across the expanse of the scaffold. Samples were placed into an incubator for two hours for initial attachment, then an additional 1000 µL of medium was added to each well. Media changes were made on Day 3 and 5, then every 2 – 3 days. The scaffold-cell constructs were cultured in static for 24 days to allow cells to bridge across pores. On Day 24, the constructs (N = 18) were moved into the stainless steel 9-well bioreactor plates (two plates for dynamic, one for static) in 550 µL of fresh media. After 24 hours to acclimatise to the new plates, constructs were mechanically stimulated by cyclic stretching to 4% sample strain (600 µm displacement) at 0.5 Hz for 4 hours a day on Day 25, 26 & 27. A preload of 1% sample strain (150 µm displacement) was applied across the 3 day loading regime. Samples were collected for analyses 2 hours after the loading cycle on Day 25, and 24 hours after the loading cycle on Day 27.

### Metabolic activity

Metabolic activity was analysed by moving constructs (Dynamic N=3, Static N=1) to fresh 6-well plates and incubating in AlamarBlue® assay reagent. Activation of the AlamarBlue® reagent was measured using a ClarioStar Plus plate reader (BMG Labtech, Germany) with an excitation wavelength of 545 nm and emission wavelength of 590 nm. Values were reported as a mean of six-readings from each sample, then the means compared using two-way ANOVA.

### DNA concentration

The DNA concentration of constructs at each timepoint were determined using the Quant-iT PicoGreen dsDNA quantification assay (Invitrogen, USA). Constructs from removed from the dynamic (N=3) and static (N=1) bioreactor plates, weighed, and frozen on Day 25 and 28. The constructs were cut into equal halves perpendicular to the strain axis, weighed, then one half digested overnight in phosphate-buffered EDTA (PBE) containing 0.5 mg/mL proteinase K (Invitrogen, USA) at 60 °C on a Thermomixer (Eppendorf, Germany). Constructs were completely dissolved over the digestion, resulting in clear lysate solutions. Triplicate aliquots of each lysate were combined at a 1:10 dilution with Quant-iT PicoGreen dsDNA reagent in Tris-HCl/EDTA (TE) buffer (1:200) to a total reaction volume of 200 uL, then incubated for 5 minutes in darkness. The fluorescence of each solution was then measured with an excitation wavelength of 480 nm and emission wavelength of 520 nm on a ClarioStar Plus plate reader (BMG Labtech, Germany). DNA concentration was determined by linear regression using a standard curve made from Lambda DNA standard provided in the PicoGreen assay kit.

### Fluorescence microscopy

Triplicate constructs from the dynamic and static bioreactor well plates were removed at days 25 and 28 and prepared for fluorescence microscopy. The constructs were fixed in 4% (w/v) paraformaldehyde (PFA) for 30 minutes at room temperature, washed in PBS, then permeabilised in 0.2% Triton-X (Sigma-Aldrich) in PBS for 5 minutes with shaking. The samples were then washed in PBS and blocked by incubating in 1% bovine serum albumin (BSA) for 10 minutes. The samples were then washed in PBS and stained for 45 minutes in darkness in DAPI and Alexa-Fluor 488 Phalloidin in PBS at ratios of 1:4:1000 respectively. Stained samples were stored in PBS at 4 °C until imaged. High fidelity fluorescence microscopy was performed using an Olympus FV4000 confocal microscope. Representative z-stack images of construct regions were taken using the Galvano scanning mode with 3.4 µm slice separation. Tile z-stack images of the loading area of constructs were taken using the resonant scanning mode with 5 µm slice separation. A 10x objective was used for both scan modes. Additional z-stack fluorescence images were obtained for samples loaded to 4% sample strain using the strain imaging tool described in **Section 3.1.6**. Local strain within pores was then measured from the images using the NCORR digital image correlation (DIC) MATLAB toolbox [71], [72].

### 3.3 Statistical analysis

Statistical analyses were performed using GraphPad Prism 10 Software (Dotmatics, USA). Differences between groups were determined using one and two-way analysis of variance (ANOVA) tests where appropriate, with values of P <0.05 considered significant. Assessment of statistical differences between groups were performed based on Tukey multiple comparison tests, with a confidence level of 95 % (P < 0.05) where the symbols (P < 0.001***, 0.001 < P < 0.01**, 0.01 < P < 0.05*) in figures indicate significance. Live/Dead and Alamar Blue assays were performed in triplicate and compared using two-way ANOVA.

## 4 Results

### 4.1 Validation of movement precision

For the static location accuracy test for the loading arm, the bioreactor was found to move to each position with an average error of 6.5 µm (**Figure 8A**). Within the context of stimulating 15 mm scaffolds within the OpenStrain Bioreactor well plate, the average error corresponds to 0.04 % of the total scaffold length. A linear regression of the experimental displacement over the programmed displacement is 𝑦 = 1.007𝑥 − 0.002 (R^2^ = 1.000). This is comparable to the linear regressions reported by Janvier and Banik for their devices, at 𝑦 = 0.992𝑥 − 0.0589 (R^2^ = 1) and 𝑦 = 0.2428𝑥 (R^2^ = 0.9992) respectively [5], [69]. The primary actuator within the study by Janvier is the commercial bioreactor platform EBERS-TC3 (Ebers Medical Technology, Spain) that can also be used independently for tensile strain culture, though to our knowledge it has only been used in one other original research paper [73]. The cyclic loading test indicated that the bioreactor could repeatably return to the same location, returning to the same pixel after each cycle in the video used for motion tracking (**Figure 8B**). Each pixel covers a distance of 160 µm.

**Figure 8.**
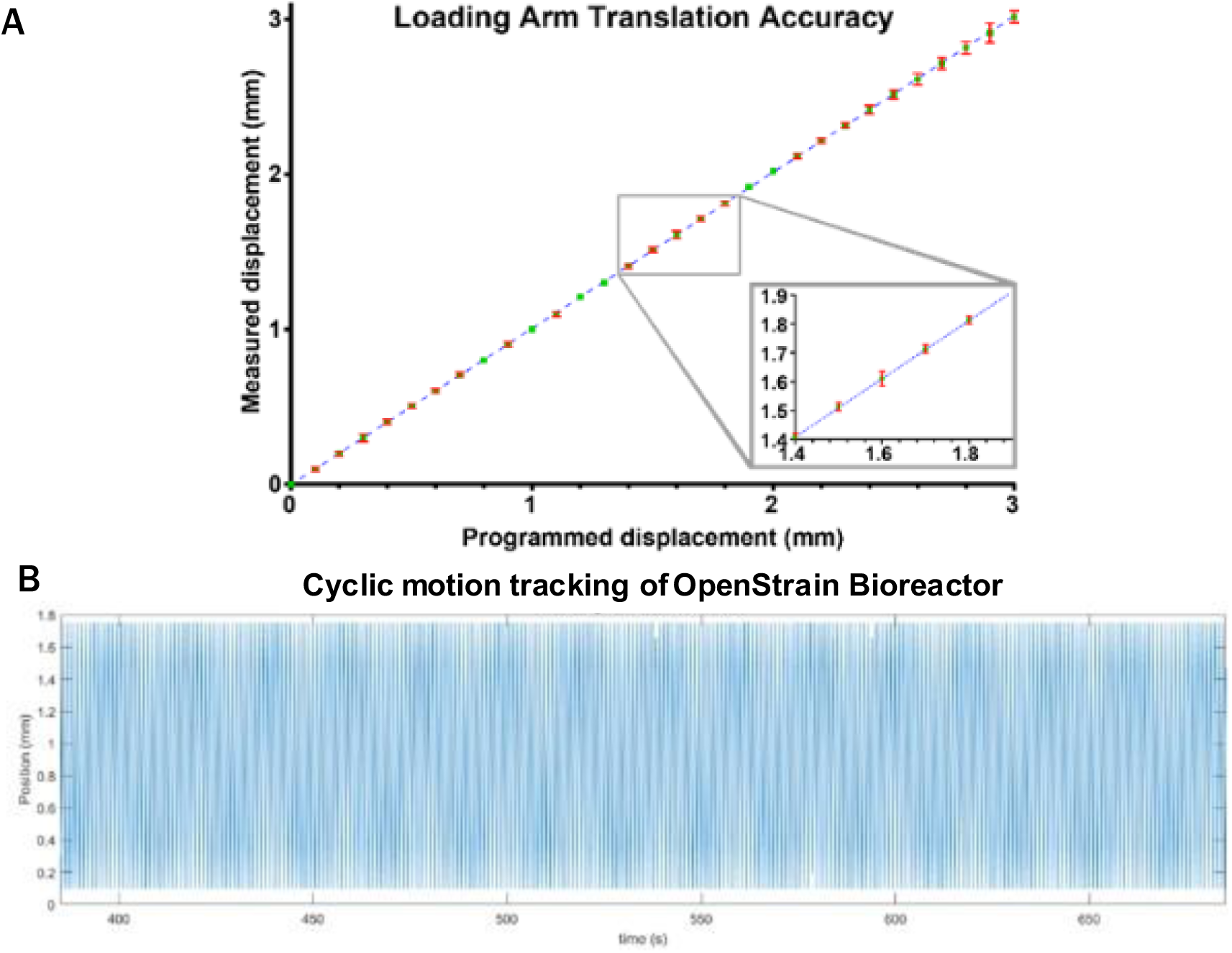
**A:** Precision of bioreactor motion in small increments between 0 and 3 mm, displaying a linear correlation between the programmed and observed displacement. **B:** Repeatability of bioreactor cyclic motion throughout a 4 hour loading regime at 1 Hz and 1.5 mm amplitude (0 to 10% strain).

### 4.2 Validation of thermal stability from stepper motors

The stepper motor began heating immediately once activated and reached a maximum temperature range of ∼ 46 °C after 120 minutes of continuous operation (**Figure 9 A**). Following the four hour loading cycle the outer surface of the stepper motor reached a maximum temperature of 46 °C. The heat of the stepper motor was dissipated by the drive box casing which did not exceed 35 °C. The heat on the outer surface of the drive box was greatest immediately adjacent to the stepper motor but dissipated through the walls of the casing reaching ambient temperature at the front of the device where the well plate attaches. The temperature of the stepper motor decreased by 3.5 °C across 10 minutes once the loading program ended, with the drive box open. The lead screw was also observed to moderately heat up over the duration of the loading cycle, approaching a maximum temperature of ∼30 °C. This heating did not noticeably extend to the lead nut. Externally visible components of the control board also reached ∼35 degrees by the 4-hour mark of the experiment.

**Figure 9.**
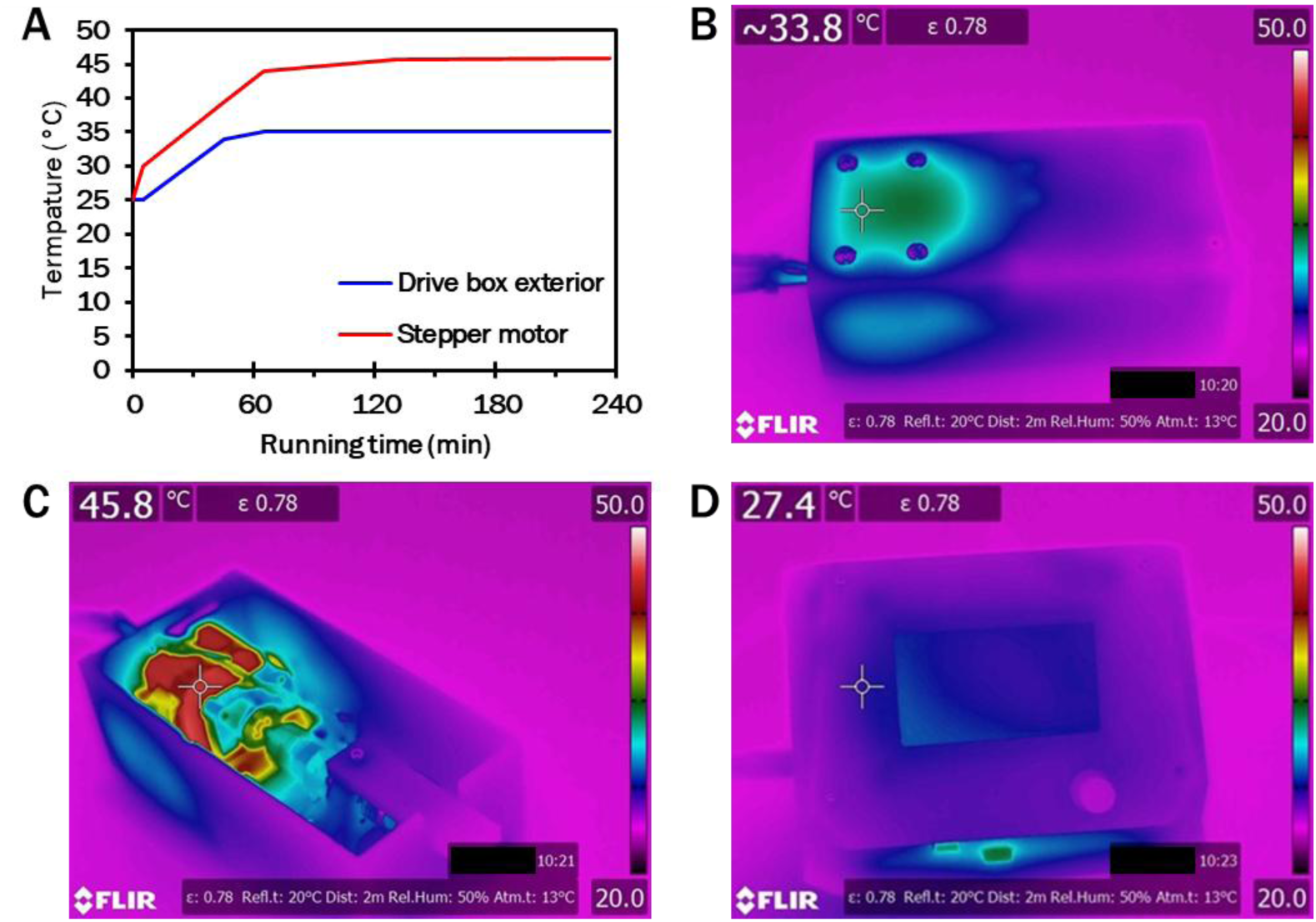
**A:** Development of stepper motor and drive box surface temperatures over four hour loading regime. **B:** Drive box surface temperature after 240 minutes of continuous operation. **C:** Stepper motor surface temperature after 240 minutes of continuous operation. **C:** Control unit casing after 240 minutes of continuous operation.

### 4.3 Static cytotoxicity study

The SSIN group seeded and cultured within the stainless-steel well plate displayed a 3.7 fold increase in metabolic activity by Alamar blue fluorescence from Day 3 to Day 7 (P < 0.05 (*))(**Figure 10 A**), with negligible dead cells visible on the scaffolds under LIVE/DEAD fluorescence (**Figure 10 B**). This is the greatest fold increase in metabolic activity over time out of all sample groups. Metabolic activity of the SSIN group was 2.4 fold higher than SSOUT seeded, though this difference was not statistically significant (P = 0.964). Constructs cultured within the stainless well plate displayed lower metabolic activity compared to constructs cultured within SBS well plates. Metabolic activity was less consistent between SS100 replicates than between SS200 replicates. The SS100 group had the highest metabolic activity of SBS well plate samples at Day 7 and the most even distribution of cell attachment on scaffolds (**Figure 10 Eii**). Cell attachment was generally uneven in all samples (**Figure 10 B-E**), with cells not reaching confluency on the surface of scaffold fibres or within pores.

**Figure 10.**
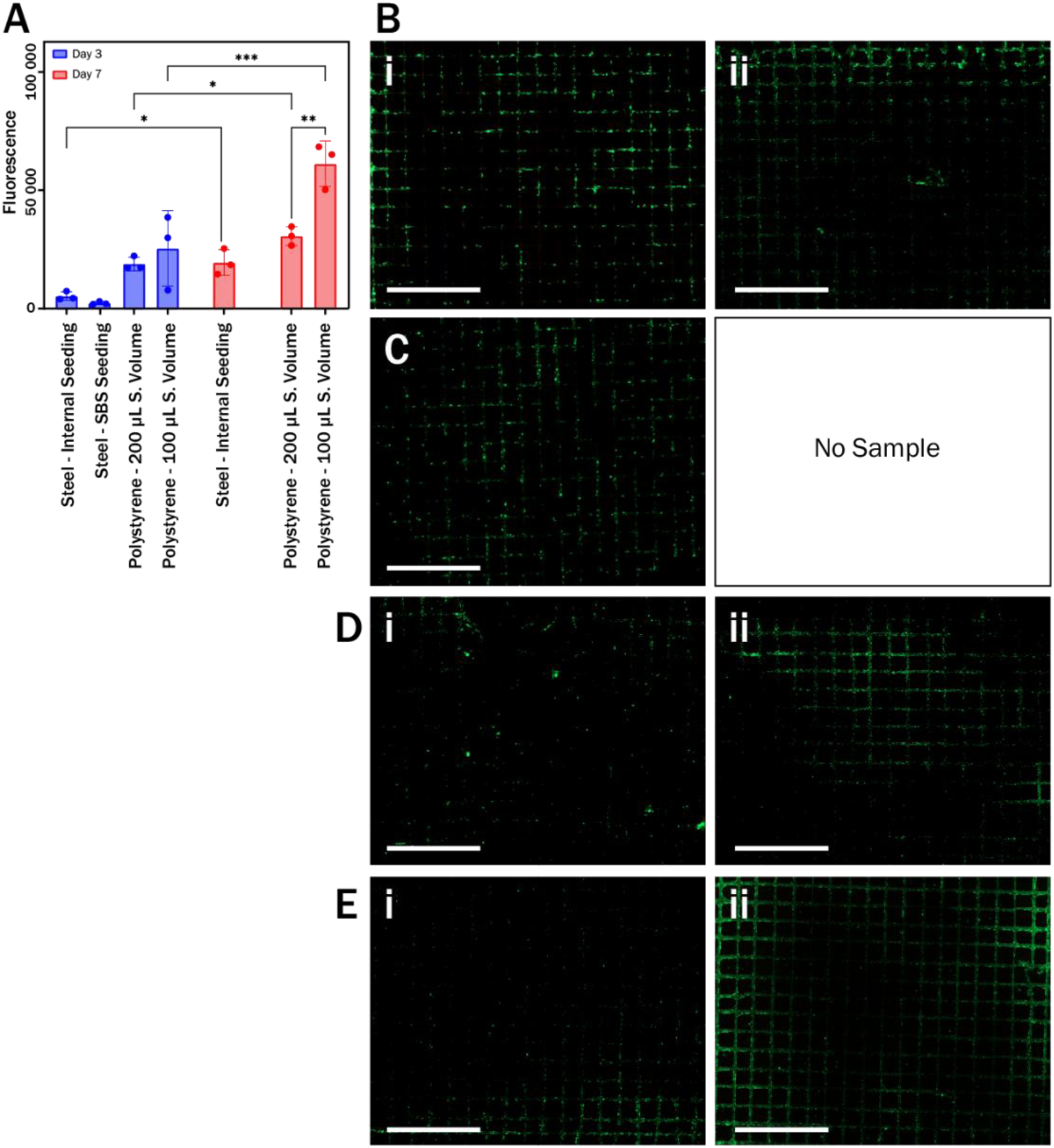
**A:** Metabolic activity of cell constructs as measured by intensity of an Alamar Blue assay stain. P < 0.05 (*), P < 0.002 (**), P < 0.001 (***). Constructs cultured within steel wells are not compared to constructs cultured within polystyrene SBS wells as the scaffold dimensions are not equivalent. **B:** Day 3 (**i**) and Day 7 (**ii**) viability of constructs cultured within the stainless-steel plate, originally seeded within the stainless-steel wells. **C:** Day 3 Viability of construct cultured within the stainless-steel plate, originally seeded within an SBS plate then moved to the stainless-steel well plate on Day 0. **D:** Day 3 (**i**) and Day 7 (**ii**) viability of construct cultured within an SBS plate and seeded with a 200 µL seeding volume. **E:** Day 3 (**i**) and Day 7 (**ii**) viability of construct cultured within an SBS plate and seeded with a 100 µL seeding volume. **B – E** All imaged under fluorescence microscopy by Calcein-AM (green, living cells)/Ethidium homodimer-1 (red, dead cells). Scale bars: 2500 µm.

### 4.4 Dynamic *in vitro* tissue construct model

#### 4.4.1 Effects on cell activity

The average metabolic activity of the dynamic samples decreased by 49% from Day 25 to Day 28 (P < 0.001***) (**Figure 11 A**). This appears comparable to the static sample (48% reduction). Construct DNA content decreased by 12% over the timeframe and appears consistent between with the static sample (20% reduction) (**Figure 11 B**).

**Figure 11.**
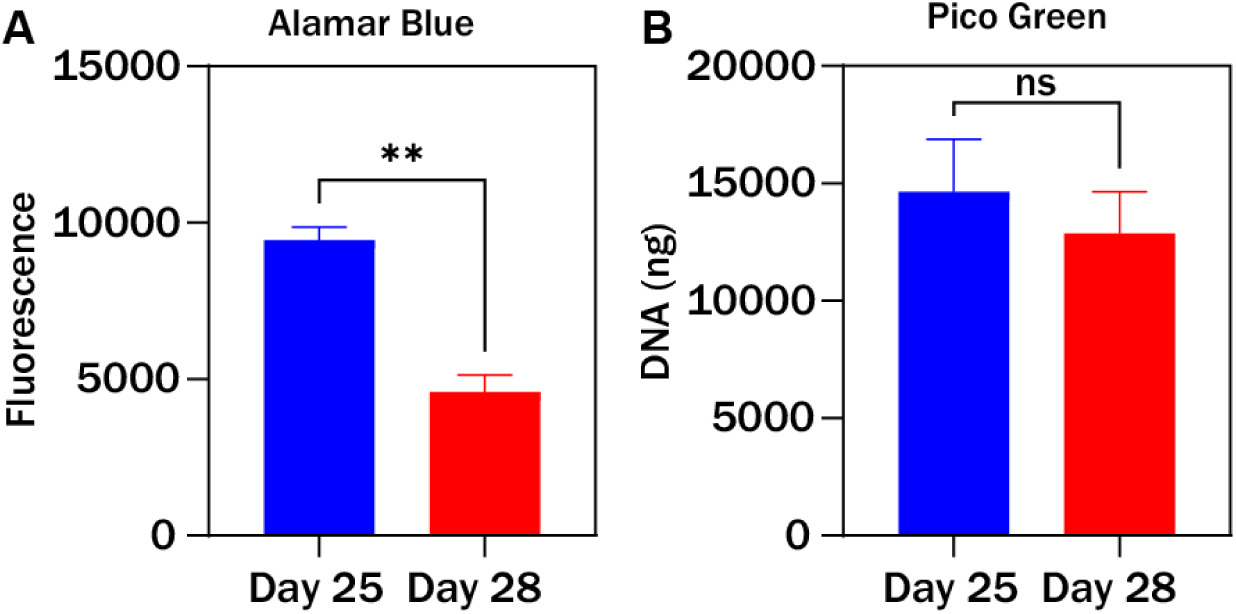
**A:** Metabolic activity of cell constructs as measured by intensity of an Alamar Blue assay stain. P < 0.05 (*), P < 0.002 (**), P < 0.001 (***). **B:** DNA concentration per tissue mass for dynamic and static constructs. P < 0.05 (*), P < 0.002 (**), P < 0.001 (***).

#### 4.4.2 Cell sheet morphology

Cells were dispersed unevenly across the scaffolds in all sample groups, ranging from pores with little to no bridging, to entirely confluent pores filled with dense tissue. Dynamic samples appear to have greater cell densities than static at equivalent time points on average, however due to the variability of scaffold cell coverage by Day 24 prior to any stimulation, it is unknown if this difference is directly caused by mechanical stimulation. Actin fibres can be found aligned with PCL fibres in all sample groups. Large regions of parallel aligned actin fibres along the uniaxial strain axis can be observed in dynamic samples at Day 28. These fibres continue across pore boundaries, at an angle of approximately 45° from the PCL fibres.

#### 4.4.3 Experimental local strain

DIC indicated substantial differences in local strain within cell sheets bridging fully confluent pores relative to on scaffold fibres (**Figure 12**). The strains within cells were reported as between 20.0% tensile strain to -19.9% compressive strain. The range of local strains in cell sheets appeared to be consistent across different areas of the scaffold.

**Figure 12.**
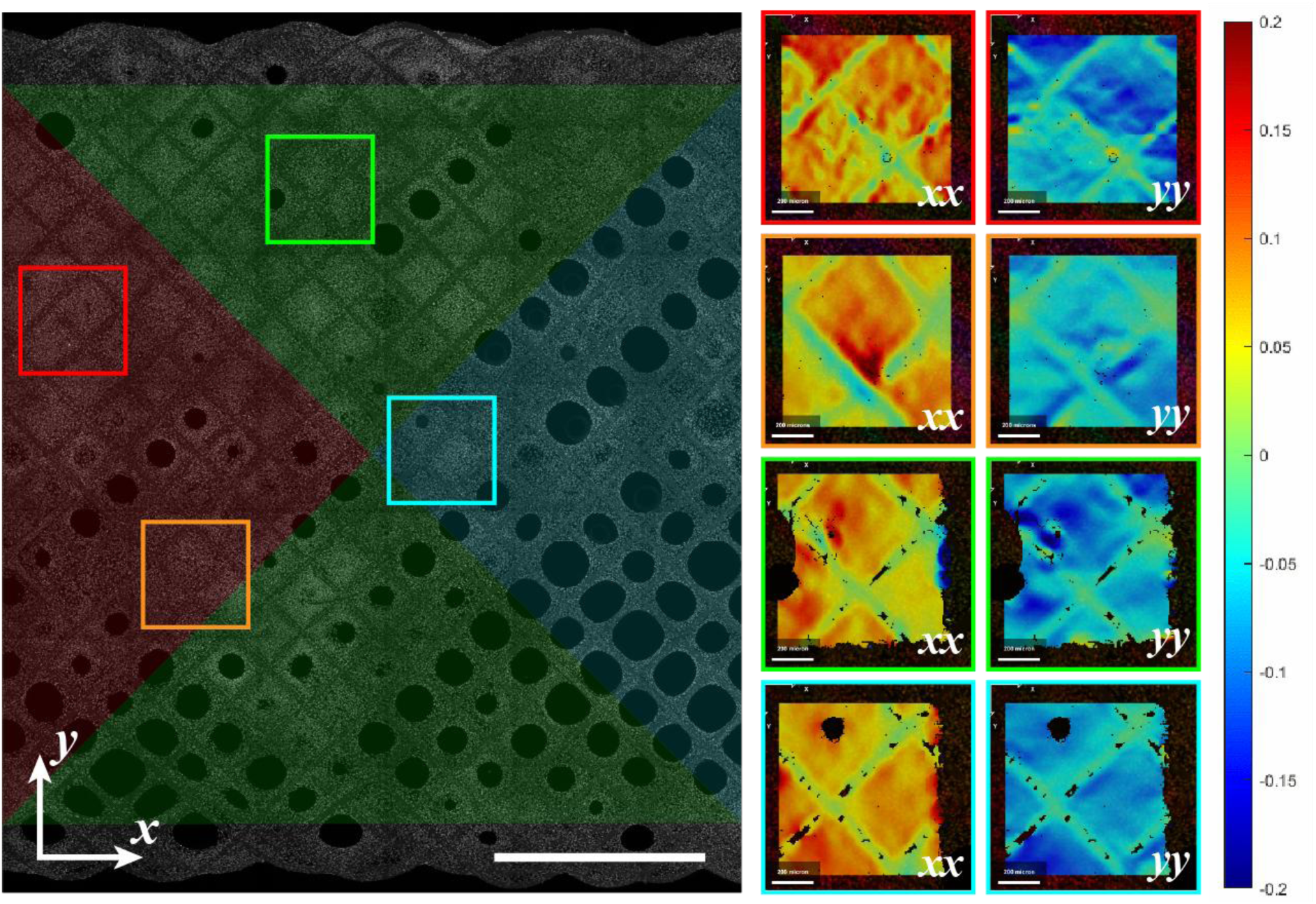
**Left:** Right half of Day 28 dynamic scaffold at 0% strain under DAPI fluorescence with simulated approximate strain levels marked in red (maximum), green (medium) and cyan (minimum). Scale bar: 2000 µm. **Right:** Local strains across the scaffold at 4% global strain determined using DIC. Scale bars: 200 µm.

**Figure 13.**
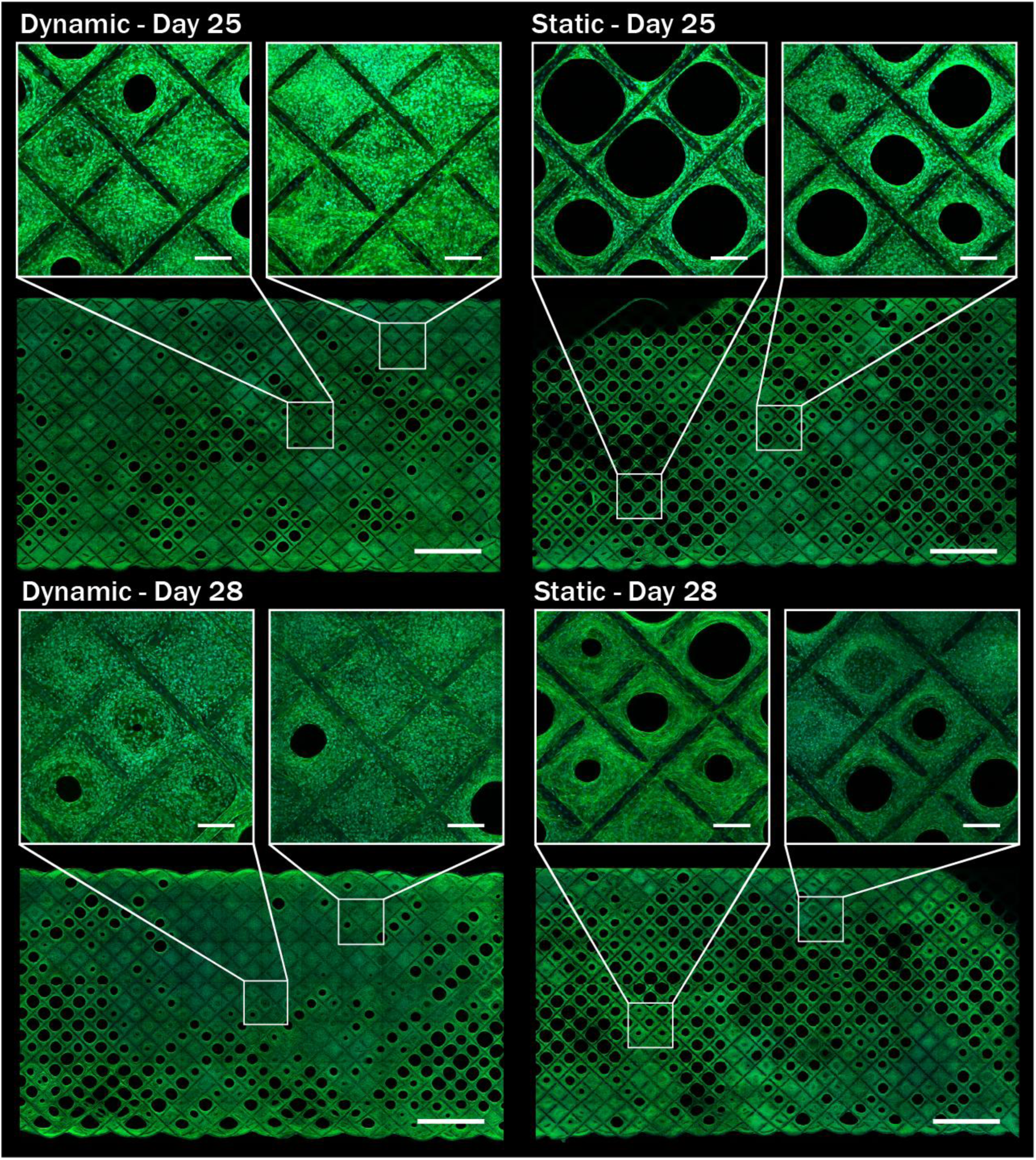
**A:** Cell morphology of sample groups by DAPI (cyan, cell nuclei)/Phalloidin (green, actin). Major scale bars: 2000 µm. Minor scale bars: 200 µm.

## 5 Discussion

### 5.1 Dynamic stimulation of cell constructs

#### 5.1.1 Cell responses

The metabolic activity and DNA concentration of constructs dropped over the three-day stimulation period. This is similar to the results obtained by Baker *et al*, who observed that applying CTS of 6% at 1Hz for 3h/day for 2 weeks to MSCs after 6 weeks of static culture resulted in a significant decrease in DNA concentration in the cell constructs compared to the unloaded constructs [55]. A further 2 weeks of CTS did not further affect DNA content, though collagen content was increased. These contrasting responses of the cell constructs to mechanical stimulation may correspond to the maturity of the cell populations and the differences in available space for cells to grow into in the constructs.

Mechanostimulation having the greatest impact shortly after seeding and reducing over longer durations is a familiar effect within the literature. As seen by Jiang *et al*, under cyclic tensile strain (CTS) the osteogenic differentiation of human periodontal ligament cell monolayers reach their highest Runx2, alkaline phosphatase (ALP) and osteocalcin (OCN) expression after 24 hours of 5% cyclic sample strain before then diminishing after an additional 24 hours of stimulation to a level more similar to after 6 to 12 hours of stimulation [53]. For bone marrow stromal cells, CTS of 5% significantly increased ALP production after 24 hours, and this level was maintained over an additional 24 hours of stimulation [39]. For groups exposed to 10 or 15% strain however, ALP production was significantly reduced by the additional 24 hours of stimulation. Haasper *et al* found that when straining bone marrow derived MSCs to 8% and 2% for three days as cell monolayers, both strain groups reached their maximum Runx2 expression by the end of the first day, before linearly decreasing over the following two days until they were indistinguishable from the controls after the third and second day respectively [32]. Rat MCS stimulated with 0.2% CTS similarly led to upregulated ALP activity 6 hours into stimulation, but this diminished back to its initial level after 24 hours [35]. The gradual desensitising of bone cell types to mechanical stimulation in bioreactor studies may be tied to a cell level expression of Frost’s mechanostat, with the cells adjusting their morphology to suit the applied mechanical environment before bringing their responses back to an initial set point [14], [15].

#### 5.1.2 Strain distributions within cell constructs

Cells within densely bridged pores experienced notably different local strains when the PCL fibres bordering the pores were displaced by the strain applied to the scaffold as a whole. This behaviour has been recognised for 2D monolayer-based tensile strain bioreactors [74], but is rarely considered for 3D cell constructs undergoing tensile loading, or is specifically assumed to be constant throughout the depth of the construct [47]. This assumption was not true for the constructs and loading conditions within this work. The 4% global tensile strain applied to the scaffold resulted in local strains within the cell populations ranging from 20.0 % tensile strain to -19.9% compressive strain. This highlights the important of considering strains at scales relevant to cells rather than only with respect to entire constructs. The strain experienced by cells in dense confluent pores will be quite different to cells in thinner bridged pore sheets, and different again to cells bordering open pores. This complicates directly linking results tied to the entire construct (ie metabolic activity from Alamar blue, or osteogenic effect through ALP activity) to specific strains, but this could possibly be achieved through image-based techniques such as immunofluorescence staining, in combination with DIC for quantifying local strains.

### 5.2 User-centred design

A particular focus throughout the development of the OpenStrain Bioreactor was to make it as user-friendly as possible. “User-friendly” is a relatively vague measure, it was approached for example by changes that decrease the number of steps required to perform certain functions with the system (such as loading scaffolds), or that decrease the required dexterity of the user to use the system, or that reduce the likelihood of risks associated with using the device. For instance, while earlier functional designs of the bioreactor mechanical unit had the drive box simply sitting on the base plate, it design was updated to include a locking system to secure the drive box to the base plate which enabled the entire assembly to be picked up with one hand, from any angle, without risk of overturning the well plate. Similarly, the control display uses a colour touch screen rather than a simpler text-based LCD screen as is found on many 3D printers, a change that makes the interface easier to navigate and use. It additionally provided the ability to change actuation speed live through a simple user interface and monitor the duration of programs.

The arrangement of the well plate and clamps is intended to minimise the required dexterity for using the device, and the complexity of steps required to prepare samples for and in the device. The clamps, for example, have dedicated tabs for manipulating them with forceps. We expect over many uses this design feature will lead to reduced likelihood of experiments failing when compared to other devices of similar function within the literature and commercial market. The CellScale MCT6 for example uses small M2 screws in deep wells, to fasten minimally featured plates (**Supplementary Figure 1**). These plates cannot be easily handled with forceps, and the small scale of the screws and high number of them increase the likelihood of dropping them within the wells during handling. The stretching device produced by Lin *et al* similar suffers from deep clamps that frustrate careful positioning of samples [61]. Other devices require extremely specific forms of samples to be compatible with their attachment device, such as samples with regions that can be embedded in epoxy [59] or bonded by infiltration into coral or bone anchors [60]. Screw-fastened clamps are a more versatile approach, and they are used in many bespoke research bioreactors [55], [56], [61]–[63], [65]–[67], [75].

When used with its maximum of five bioreactor units, the OpenStrain Bioreactor system can stimulate 45 samples simultaneously and each bioreactor unit can be given a unique program. To our knowledge, this is the greatest number of individual samples for any singular tensile bioreactor system for 3D samples out of both commercial and bespoke research bioreactors within the context of applying tensile strain (**Table 3**). The device with the next highest throughput is the Ebers TC-3 with a 20-well attachment. Even these well numbers still pale in comparison to typical SBS plates however, so it is desirable to redesign the well plates to be wider and contain more wells. With the limitation of keeping the well plate within the dimensions of an SBS plate, wells would need to be thinned for more to fit. If this limitation is loosed however then the plates can more easily be expanded. More powerful stepper motors may be required if the number of wells is increased substantially.

**Table 3.** Summary of number of wells per bioreactor for commercial tensile strain platforms compared to the novel OpenStrain Bioreactor and estimated price per well.

| Bioreactor | Number of wells | Well volume (mL) | Purchase cost (AUD) | Cost per well (AUD) |
| --- | --- | --- | --- | --- |
| OpenStrain Bioreactor (minimum well number) | 9 | 0.7 | \$2,000 | \$222 |
| OpenStrain Bioreactor (maximum well number) | 45 | 0.7 | \$6,400 | \$142 |
| Ebers TC-3F (standard) | 3 | 90 | Unknown | Unknown |
| Ebers TC-3F (high throughput) | 20 | 5 - 11 | Unknown | Unknown |
| CellScale MCT6 | 6 | 100 - 300 | \$17,000 | \$2,833 |
| CellScale MCJ1 | 6 | 30 - 150 | \$50,100 | \$8,350 |
| TA Instruments Electroforce BioDynamic 5500 | 4 | 102 - 212 | \$77,000 | \$19,250 |

The design of the bioreactor system was strongly influenced by its intention to be easily fabricated by researchers with little manufacturing experience, ultimately at similar difficulty level to assembling a DIY 3D printer or hobby electronics kit. The stainless steel parts are the most difficult to manufacture as they require the use of a CNC mill which may not be available for research groups in house, however custom CNC machined parts can now be cheaply produced through external machining companies such as SunPE or PCBway. If the bioreactor was to be produced at a higher throughput for commercialisation there are several changes recommended for improving the design and manufacturing processes. To minimise the cost-per-unit for the well plate lids it would be more suitable to fabricate the plates at high-throughput from polystyrene using injection moulding. This is how the base structure of standard SBS well plates are made. Injection moulding would also be an ideal tool for manufacturing the bioreactor casing at lower unit-cost and greater speed than 3D printing. Remaking the control board to use a custom PCB designed specifically for the requirements of the bioreactor would also be beneficial, and more robust than using a retrofitted 3D printer control board. Use of a custom board may allow for additional functionality not currently possible with the current board, such as live datalogging from force sensors, or additional stimulation modes such as electrical or ultrasonic. Maintaining the design as open-source would also allow for users around the globe to design new functionalities for the system and substantially increase its applications.

### 5.3 Design for sterilisation

Minimising the risk of contamination is at the forefront of any cell culture experiment, with researchers constantly adhering to the most stringent of protocols. Standard cell culture tools like serological pipettes or SBS well plates have well established and research methods for sterilisation and sterile use, and it’s vitally important for new devices to aim to match these. There is a tension in this however as often changes that make a device more complex or versatile also increase the difficulty of reliably decontaminating and sterilising them. Consider the differences between an orthopaedic manual hand saw and a battery powered bone saw as an example. Unlike the manual hand saw, the powered bone saw can’t be simply put into an autoclave, or entirely disassembled and individually sterilised as parts, or entirely disposed of and replaced after each use. When designing sterilisable mechatronic devices, the designers must determine an acceptable target of what is ‘sterilisable enough’. It’s similar in concept to designing a building for a particular Factor of Safety, or to resist a certain frequency of natural disaster like a ‘one in a hundred year earthquake’. For designing the OpenStrain Bioreactor, the discussion around decontamination and sterilisation as most significant for the well plate assembly. The simplest method of securing scaffolds in the well plate that we could conceive was to pierce them on spikes. Seemingly without the need for screw threads, this would be the most easily cleanable securement method, but in prototypes it wasn’t sufficiently reliable for securing scaffolds, especially over many cyclic loads. The concession was to use screw tightened clamps that could hold scaffolds reliably. This did introduce screw threads that would be more difficult to clean than smooth spikes, though it was at least ensured that the threads would be located as far as reasonably possible from the media solutions within the wells. This is similar in principle to the approaches used by Leung, Baker and others in their bespoke bioreactor systems that secure samples with screws immediately proximal to but not within media volumes [32], [55], [59], [61].

We expect that over many uses, there will ultimately be fewer contaminations in wells resulting from insufficiently decontaminated screw threads than other bioreactor designs that use screw threads directly within media volumes, such as the CellScale MCT6 and MCJ1 or bespoke research bioreactors such as by Raimondi, Qin, Corti, and numerous others [56], [62], [63], [65], [66], [68], [75], [76]. A concession made for the sterilisation of the bioreactor drive box was to fabricate the parts using FDM 3D printing PLA. The layer lines and porous nature of the material undoubtedly present more of a risk of insufficient decontamination between tests, however as the parts are located far from the sealed well plate and due to the limited time and budget available to make alternatives, they were deemed sufficient. With more time and budget available, and to make a greater number of devices, it is recommended that the design of the drive box be reconsidered to use materials more common for cell culture tools, such as injection moulded polypropylene or folded stainless-steel.

### 5.4 Scaffold fixation – an open debate

The method to securely hold constructs within the bioreactor presented significant challenges in the design process. The problem of how best to secure biological samples for tensile tests and loadings is widely investigated and has no single solution. Clamps are the most common method as they can ensure samples are loaded uniformly at each clamped end, though despite their frequency they still introduce challenges [57]. As biological samples are typically soft with high water content they can slip out from smooth clamps, and if the clamping pressure is increased the samples can be damaged from the deformation before the pressure is sufficient to secure them [61], [77], [78]. The clamps also introduce concentrated shear stress zones at their surfaces that do not extend the full depth of samples and may interfere with loading or induce failures within the gauge region [79]. Pin-based securement systems may more uniformly load the entire depth of samples and can prevent slipping, though they require constructs either custom designed for the pins or that samples be damaged by the pins and they can also create undesirable stress concentrations [58], [59], [80]. Adding serrations, matched hill-and-valley, or self-tightening wedge features to the mating surfaces of clamps can reduce the likelihood of samples slipping though it also introduces stress concentrations which can lead to samples failing at the clamp edges [81], [82]. Concessions for stress concentrations, ease of handling and manufacturing complexity must all be balanced based on the particular samples under investigation. For the current project screw-fastened clamps with a hill-and-valley ridge were sufficient for the MEW scaffolds investigated. To improve their versatility further a combined pin-and-clamp approach (**Supplementary Figure 2**) alike to the proposed clamp design by Jiang may enable softer samples such as hydrogel constructs to be strained using the OpenStrain Bioreactor [57]. A tool for simultaneously loading multiple clamps would also benefit cell culture experiments using the device by reducing the time for loading samples, especially samples containing live cell populations.

### 5.5 Investigation of cytotoxicity

It is well known that 316 stainless-steel is biocompatible, it is one of the most common materials for metal implants. As the stainless-steel parts in this study were made by a more generalised manufacturing service not specialised in producing medical devices however, the cytotoxicity study confirmed that the parts had not been made cytotoxic during the manufacturing process. It also confirmed that the acrylic well plate lid formed a suitable closed environment akin to the lid on an SBS well plate. These were necessary confirmations prior to the use of the parts in live dynamic cell studies. The stainless-steel parts used for this study were manufactured via DeFab by a third-party online manufacturing service, SunPE, that could be used by other research groups for their own custom machining. Services such as SunPE or PCBway have made designing advanced parts from custom materials more accessible, as it means research groups aren’t limited to just the manufacturing equipment and materials they have internally.

## 6 Conclusions

A custom bioreactor for culturing cell-seeded scaffolds under cyclic tensile strain over prolonged *in vitro* culture was designed using an iterative design process. The system, referred to as the ‘OpenStrain Bioreactor’ applies strain to cell constructs through a displacement-control method. The device was evaluated with respect to its accuracy and precision of movement, cytotoxicity, and ease of use, and deemed to be appropriate for use in mechanically dynamic cell culture. It was used to apply cyclic tensile strain to MC3T3 osteoblasts on PCL MEW scaffolds over 3 days in a 28 day *in vitro* experiment. Due to the diamond shaped repeating unit cell for the scaffolds, a wide range of local strains were applied to cells depending on their locations within the scaffolds. In comparison to existing commercial tensile strain platforms and published experimental bioreactors, the OpenStrain Bioreactor has advantages in well number, device footprint, assembly cost, and ease of use. The OpenStrain Bioreactor has acceptable precision and accuracy to be used for imparting cyclic tensile loading to scaffolds for *in vitro* experiments. In these it is comparable to other similar devices within the literature, including commercial bioreactors. The stainless-steel well plate has acceptable cytotoxicity for use within *in vitro* cell culture experiments. The design for the device will be released to the public as open-source to encourage collaborative research and repeatability between tensile strain experiments of disparate groups. It is believed that the OpenStrain Bioreactor will be useful for a wide range of tensile strain mechanoculture experiments both for bone cells and for other biological cell types such as tendons or cardiac tissue, or more [2], [6]–[11].

