## Supplementary Figures for "Development of a displacement-controlled uniaxial-strain bioreactor for high-throughput, dynamic *in vitro* cell culture"

**Supplementary Figures 1-2**

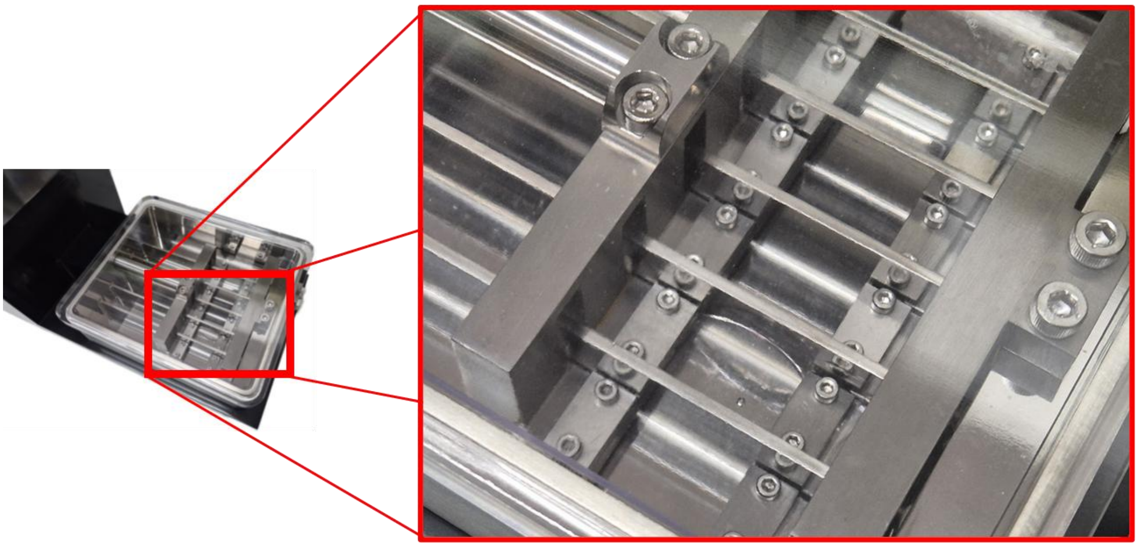

**S. Fig. 1:** Fixation system of CellScale MCT6 commercial bioreactor with multi-well configuration without media solution. A length of plastic is fixed within the second well using the clamping plates and M2 metric machine screws.

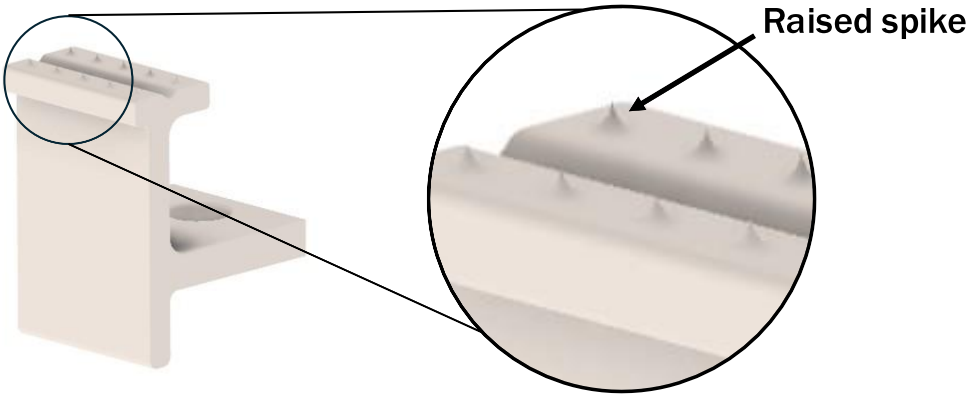


**S. Fig. 2:** Potential alternative clamp design derived from OpenStrain clamp design, featuring 10, 0.5 mm tall 3D printable raised spikes (circled) on the mating surface for securing hydrogel samples. Clamp has a total height of 15.5 mm including the spikes.
